# PARIETAL CONTRIBUTIONS TO ABSTRACT NUMEROSITY MEASURED WITH STEADY STATE VISUAL EVOKED POTENTIALS

**DOI:** 10.1101/2020.08.06.239889

**Authors:** Peter J. Kohler, Elham Barzegaran, Anthony M. Norcia, Bruce D. McCandliss

## Abstract

Non-symbolic number changes produce transient Event Related Potentials over parietal electrodes, while numerosity effects measured with Steady-State Visual Evoked Potentials (SSVEPs) appear to originate in occipital cortex. We hypothesized that the stimulation rates used in previous SSVEP studies may be too rapid to drive parietal numerosity mechanisms. Here we recorded SSVEPs and behavioral reports over a slower range of temporal frequencies than previously used. Isoluminant dot stimuli updated at a consistent “carrier” frequency (3-6 Hz) while periodic changes in numerosity (e.g. 8→5) formed an even slower “oddball” frequency (0.5-1 Hz). Each numerosity oddball condition had a matched control condition where the number of dots did not change. Carrier frequencies induced SSVEPs with midline occipital topographies that did not differentiate the presence or absence of numerosity oddballs. By contrast, SSVEPs at oddball frequencies had parietal topographies and responded more strongly when oddballs were present. Consistent with our hypothesis, numerosity effects were stronger at slower stimulation rates. In a second study, the numerosity change was either supra-threshold (e.g. 8→5 dots) or near the threshold required for detecting numerosity changes (e.g. 8→9 dots). We found robust parietal responses for the supra-threshold case only, indicating a *numerical distance effect*. A third study replicated the parietal oddball SSVEP effect across four distinct suprathreshold numerosity-change conditions and showed that number change direction does not influence the effect. These findings show that SSVEP oddball paradigms can probe parietal computations of abstract numerosity, and may provide a rapid, portable approach to quantifying number sense within educational settings.

## Introduction

Numerical cognition encompasses a range of abilities including explicit computations such as addition and subtraction, counting, the appreciation of spatial constructs such as the number line as well as the ability to estimate numerical magnitude. Numerical magnitude estimation is frequently studied with non-symbolic visual tokens that vary in their numerosity or “cardinality”, e.g. the specific number of tokens in a group. The perception of numerosity and cardinality has been argued to involve separate processes, depending on the range of magnitudes involved, forming three general regimes of magnitude. Enumerating small numbers of tokens or “subitizing” has been proposed to involve parallel individuation, or “subitizing”, up to a limit of approximately 4 (Jevons, 1871; Kaufman et al., 1949; Mandler and Shebo, 1982). An “Approximate Number System” in which estimated magnitudes scale logarithmically with number of tokens has been demonstrated for quantities in a second regime between 5 and ~50 (Atkinson et al., 1976; Feigenson et al., 2004; Piazza et al., 2007; Anobile et al., 2014). A third regime has been demonstrated that invokes texture-like processing with even larger quantities (Anobile et al., 2016; Portley and Durgin, 2019).

Here we focus on neural responses generated in the approximate number range. Such responses to number were first recorded in macaque parietal cortex where neurons tuned to number were found (Sawamura et al., 2002; Nieder and Miller, 2004). Later work (Roitman et al., 2007) measured single-unit activity as a function of numerosity in the macaque lateral intra-parietal area (LIP) and found modulation of activity associated with numerosity as early as 100 msec after stimulus presentation. To accurately compare latencies across species, previous studies have derived a 5/3 species conversion factor (Schroeder et al., 1995; Chen et al., 2006). LIP responses to number were independent of attention, reward and a number of low-level stimulus attributes such as size, color or density. Studies using functional MRI (fMRI) in humans have also implicated parietal cortex as central to approximate number estimation, identifying multiple, topographically organized maps of numerosity in parietal cortex starting in the horizontal extent of the intra-parietal sulcus (hIPS) and extending to the post-central sulcus, with an additional map in temporal occipital cortex (Piazza et al., 2004; Harvey et al., 2013; Harvey et al., 2015; Harvey and Dumoulin, 2017).

Neural responses in the approximate number range have also been measured with transient event-related potentials (ERPs). ERPs that were sensitive to numerical difference during the performance of non-symbolic comparison tasks have been found over parietal electrodes at latencies around 200 msec (Temple and Posner, 1998; Libertus et al., 2007; Heine et al., 2011; Heine et al., 2013). ERPs sensitive to adaptation to non-symbolic numerical quantity have also been found in parietal regions using source estimation (Hyde and Spelke, 2012).

The neural coding of non-symbolic numerosity has also been studied with steady-state visual evoked potentials (SSVEPs). By contrast with the transient ERP studies reviewed above, these studies have found their dominant response at electrodes over early visual cortex. Park and colleagues (2018) updated dot cloud images at 8 Hz and imposed additional periodicity at 1 Hz by updating numerosity and, in separate runs, other dimensions such as size, density, total area. The 1 Hz response was largest for the numerosity condition and was maximal over posterior midline electrodes near Oz and right hemisphere electrode PO8. The number-related responses were measured in an implicit task and participants were unaware of the experimental manipulation. The author concluded that numerosity is directly perceived as a primary stimulus dimension. Work from other groups (Guillaume et al., 2018; Van Rinsveld et al., 2020) has applied a more rapid periodic visual stimulation protocol in which dot clouds with a base numerosity were presented at 10 Hz with every 8^th^ cloud being replaced with a different numerosity, so that numerosity was updated at 1.25 Hz. Oddball responses grew larger as numerical distance increased and the magnitude of the oddball response was correlated with Weber fractions measured for individual participants in a numerosity judgement task. This was true even though the SSVEP stimuli were task irrelevant (Guillaume et al., 2018). In both studies, the oddball response was centered on electrodes over early visual cortex. The topography of the response suggested that numerical magnitude is automatically extracted very early in the visual system (Guillaume et al., 2018). A similar topography was observed when rapidly alternating dot clouds of unequal number were used (Lucero et al., 2020). Computational modeling (Van Rinsveld et al., 2020) has shown that early visual cortex could signal the presence of a number oddball or number alternation independently of other correlated stimulus dimensions.

The SSVEP and transient ERP studies differ in two main respects that may help explain the contrasting results. One is the temporal frequency of the stimuli, being high for the SSVEP and effectively low for the ERP, the other is whether number is task-relevant or task irrelevant. We hypothesized that parietal number mechanisms have a limited temporal acuity and that SSVEP responses over parietal cortex could be recorded using task-relevant stimuli presented at slower rates. By manipulating the SSVEP stimulation rate, we find that behavioral discrimination of numerosity is better for lower temporal frequencies and that stimulation at lower temporal frequencies gives rise to consistent activation centered over parietal, rather than occipital cortex, consistent with parietal number mechanisms having low temporal resolution and being important for conscious perception of number.

## Materials and Methods

### Participants

15 participants (7 females, mean age 27.9±8.7) took part in Experiment 1. 15 participants (7 females, mean age 20.9±4.5) took part in Experiment 2. 15 participants (11 females, mean age 24.1±8.1) took part in Experiment 3. Three participants took part in two experiments, including one of the authors, but none took part in all three. All were pre-screened to confirm that they had normal or corrected-to-normal visual acuity on the Bailey-Lovie chart and normal stereopsis on the RandDot test (http://precision-vision.com/products/stereo-vision-tests/randot-stereo-test.html). Their informed consent was obtained before the experiment under a protocol that was approved by the Institutional Review Board of Stanford University.

### Stimulus generation

Dot coordinates and sizes were generated using a previously published algorithm (Piazza et al., 2004) that was designed to ensure that standard and oddball updates were statistically matched in terms of non-number properties. Dots were drawn with a dark inner dot surrounded by a brighter outer ring of equal area to eliminate confounding luminance and numerosity (Carlson et al., 1984; Huk and Durgin, 1996; Portley and Durgin, 2019). The inner dot and outer ring were made to have approximately the same number of pixels, but rounding error made a perfect luminance match with the background impossible. To account for this, the luminance values were adjusted up or down for the outer ring, with the inner dot being kept at lowest possible luminance, such that the average luminance across outer and inner was always matched to the background luminance. Examples of the stimuli are shown in Figure 1.

**Figure 1:**
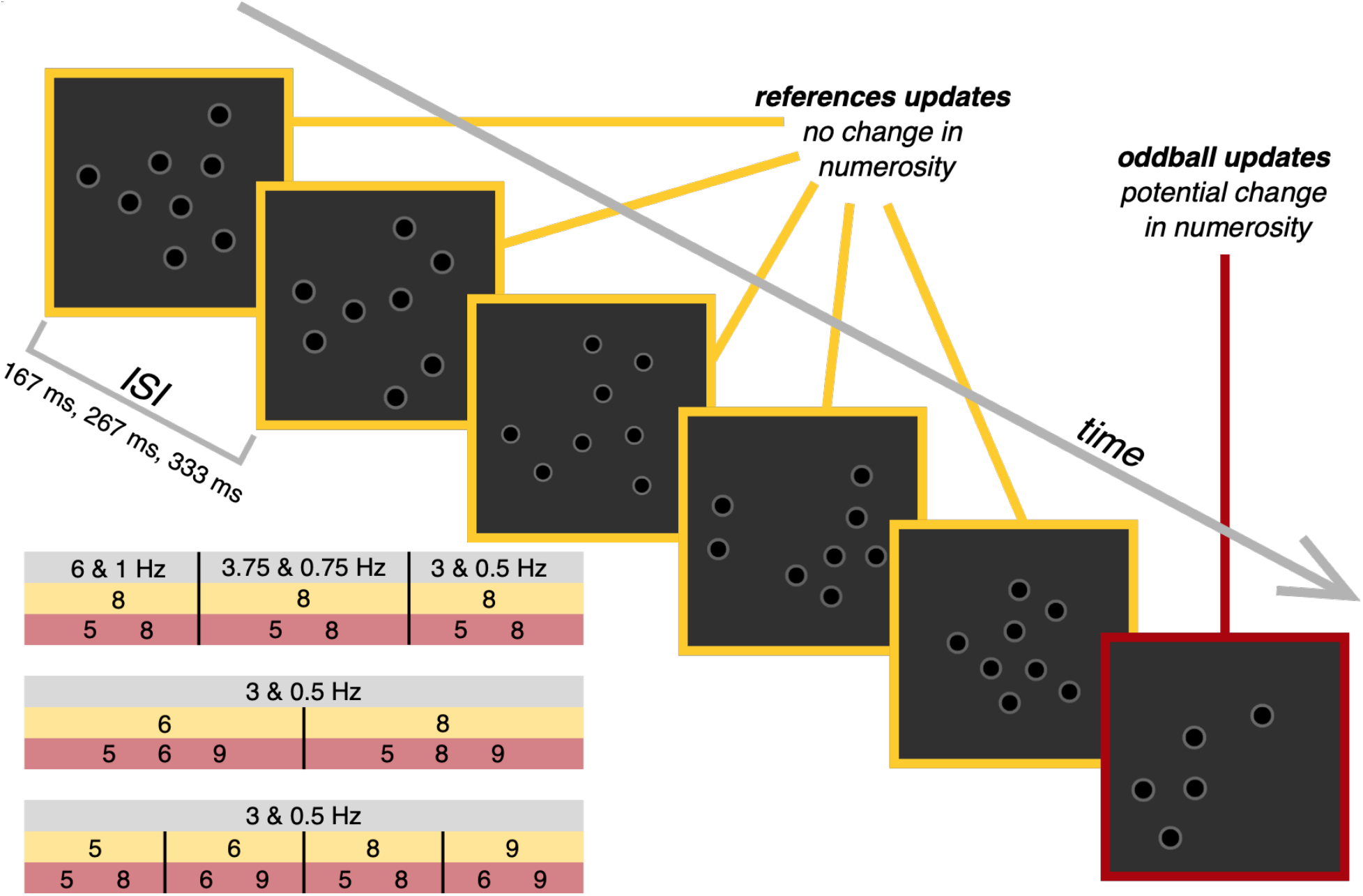
Experiment Design. Schematic showing a single cycle of the experiment, with 8 as the reference and 5 as the oddball. In Experiment 1, the carrier frequency was either 6, 3.75 or 3 Hz (corresponding to an ISI between images of 167 ms, 267 ms or 333 ms), and the oddball frequency was either 1, 0.75 or 0.5 Hz. Experiments 2 and 3 used the same frequency pairing across all conditions (3 and 0.5 Hz) but varied the reference and oddball numerosities as indicated in the table in the lower left of the figure. Each experiments included a control condition in which the oddball numerosity was the same as the reference, so that the “oddball updates” introduced no change in numerosity.

### Stimulus Presentation

The stimuli were shown on a 24.5” Sony Trimaster EL PVM-2541 organic light emitting diode (OLED) display, with a screen resolution of 1920 × 1080 pixels, 8-bit color depth and a refresh rate of 60 Hz, viewed at a distance of 70 cm. The dot images were presented in the center of the screen covering 10° × 10° of visual angle, while the rest of the screen was matched to the dot image background luminance, which was also average luminance across the whole screen. Contrast was set at 95%.

### EEG Experimental Procedure

Across our three experiments, we varied two aspects of the stimulus presentation. Experiment 1 varied the carrier and oddball frequencies, which could be one of three pairings: 6 Hz carrier with 1 Hz oddball, 3.75 Hz carrier with 0.75 Hz oddball and 3 Hz carrier with 0.5 Hz oddball. The ratio of oddball to carrier updates per cycle was in the same range for all three pairings: 1/6, 1/5 and 1/6. In Experiments 2 and 3, the slow frequency pair (3 & 0.5 Hz) was used for all conditions. The number of stimulus cycles shown per trial varied with stimulus frequency. For 1 & 6 Hz, a trial consisted of 10 cycles of 1 sec duration, for 0.75 & 3.75 Hz, 8 cycles of 1.33 sec duration and for 0.5 & 3 Hz, 5 cycles of 2 sec duration. An additional stimulus cycle was added at the beginning and end of each trial, but not included in the data analysis. This was done to reduce the influence of transient responses associated with the first appearance of the stimuli, and because participants might blink more in the beginning and towards the end of a trial. This meant that regardless of frequency pair, the total duration of each trial was ~12 sec, with ~10 sec of data going into the analysis.

Second, the reference and oddball numbers could vary across a range of numerosity pairings, although the same carrier and oddball numerosity was used on every trial within a condition. In Experiment 1, across all three frequency pairings, the reference numerosity was always 8 and paired with an oddball that could be either 8 or 5. In Experiment 2, the reference numerosity could be 6 paired with oddballs that were either 5, 6 or 9, or 8 paired with oddballs that were either 5, 8 or 9, for a total of six conditions. Experiment 3 used eight distinct pairings of reference and oddball numerosities for the eight conditions: 5 vs 5, 5 vs 8, 6 vs 6, 6 vs 9, 8 vs 8, 8 vs 5, 9 vs 9 and 9 vs 6. These pairings meant that the distance in numerosity between the reference and oddball could either be 0 (i.e. reference = oddball, used as a control condition), 1 or 3.

### EEG Acquisition and Processing

The EEG data were collected with 128-sensor HydroCell electrode arrays (Electrical Geodesics, Eugene, OR). For all participants in Experiment 1, and a single participant in Experiment 2, we used a Electrical Geodesics NetAmp400 and accompanying NetStation 5.2.0 software, while the rest of the data were collected using a Electrical Geodesics NetAmp300 and NetStation 4.3.1 software. EEG data initially sampled at 500 Hz was resampled at 420 Hz to provide 7 samples per video frame. Digital triggers were sent from in-house stimulus presentation software and stored with the EEG recording to allow synching of the visual stimulus and EEG with millisecond precision. Recordings were exported from NetStation using a 0.3-50 Hz bandpass filter, which was applied twice to ensure that power in frequencies outside the filter range was minimized. The data were then imported into in-house signal processing software for preprocessing. Prior to preprocessing, the time series collected with Netstation 5.2.0 software were shifted backwards in time to account for a 60 ms delay that is introduced in this version of the software.

If more than 15% of samples for a given sensor exceeded a 30 μV amplitude threshold, the sensor was excluded from further analysis. Sensors that were noisier than the threshold were replaced by an averaged value from six of their nearest neighbors. The EEG data was then re-referenced to the common average of all the sensors and segmented into a number of epochs (each corresponding to exactly 1 stimulus cycle). Epochs for which more than 10% of data samples exceeded a noise threshold (30 μV) or for which any sample exceeded a peak/blink threshold (60 μV) were excluded from the analysis on a sensor-by-sensor basis. If more than 7 sensors had epochs that exceeded the peak/blink threshold, the entire epoch was rejected for all channels. This was typically the case for epochs containing artifacts, such as blinks or eye movements.

### Behavioral Data Analysis

For the behavioral analysis, we computed *d*′ and criterion (*c*) based on the z-transformed probabilities of hits (*H*) and false alarms (*F*), using the following equations (Macmillan and Creelman, 2005):

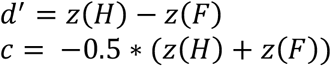

*d*′ is a measure of sensitivity and better performance (higher hit rates and lower false alarm rates) results in higher *d*′ values. Criterion (*c*) is a basic measure of bias, that will take negative values when misses are more frequent than false alarms, and positive values when the opposite relationship is observed. In the current experiments, positive values would indicate that participants have a bias towards reporting that a number change did occur, while negative values would indicate a bias towards reporting that no change in numerosity took place.

### SSVEP Data Analysis

As mentioned above, each trial consisted of several bins, with the bin length and number of bins varying depending on the stimulation frequencies (1 & 6 Hz: 10 × 1 sec; 0.75 & 3.75 Hz: 8 × 1.33 sec; 0.5 & 3 Hz: 5 × 2 sec). The amplitude and phase of the Steady-State Visual Evoked Potentials (SSVEPs) were extracted using a Recursive Least Squares adaptive filter (Tang and Norcia, 1995) with a memory length equal to the bin length. Real and imaginary components of the SSVEPs at the first four harmonics of both the oddball frequencies (referred to as *1F1*, *2F1, 3F1, 4F1*) and carrier frequencies (referred to as *1F2*, *2F2, 3F2, 4F2*) were calculated separately and averaged across bins within a trial.

We reduced the spatial dimensionality of our data by decomposing the sensor data into a set of physiologically interpretable components using Reliable Components Analysis (RCA; Dmochowski et al., 2015). Because SSVEP response phase is constant over repeated trials of the same stimulus, RCA utilizes a cross-trial covariance matrix to decompose the 128-channel montage into a smaller number of components that maximize trial-to-trial consistency through solving for a generalized eigenvalue problem. The bin-average real and imaginary values from the first four harmonics of both the oddball and carrier frequencies, from all 128 sensors, and from all trials and participants, served as the input data for RCA. For each experiment, reliable components were derived using data from all conditions, except for Experiment 1 where reliable components were derived separately for each of the three pairs of stimulation frequencies. This gave us three sets of components for Experiment 1, each based on data from two conditions, and a single set of components for Experiment 2 and for Experiment 3. To investigate the response of each component in the time domain, we also projected bin-average time domain data through the component weights that were derived based on frequency domain data. Our analyses focused on the first two RC components, which explained a substantial amount of the reliability and variance in the data, as detailed below.

After projecting the bin-average real and imaginary data through the component weights, averages across trials in each condition and across participants were generated using vector averaging, in which the real and imaginary coefficients for a given harmonic are averaged across participants, and the amplitude is computed from the result. For statistical analysis, we determined the magnitude of the projection of each participant’s vector onto the group vector average (Hou et al., 2009). The magnitudes of these projections were then used to compute the standard error for each condition, and as input for statistical testing (see below). The mean of these projected amplitudes is the same as the amplitude of the vector average. The projection procedure is useful because it preserves the robust phase consistency across subjects with associated SNR improvements that would not occur if individual participant amplitudes were used to compute errors and conduct statistical testing. For the time-domain data, the mean and standard error across participants were computed at each time-point.

### Statistical testing

We ran a linear mixed-effects analysis (LMEA) separately for data from each component, separately for harmonics of the oddball and carrier frequency, and separately for each stimulation frequency and reference number. Harmonic (*1F1*, *2F1, 3F1, 4F1* or *1F2*, *2F2, 3F2, 4F2)* and distance between reference and oddball number (0, 3 or 0, 1, 3) were within-participant factors, with participants as a random effect. The LMEAs were implemented using the lme4 (Baayen et al., 2008) package in R (R Core Team, 2014) and we used an analysis of variance (ANOVA) to test for significance, implemented using the lmerTest (Kuznetsova et al., 2013) package. Our model tested for main effects of harmonic and distance, as well as for the interaction between harmonic and distance. To test the significance of the interaction, we also ran a second model that left out the interaction term, and ran an ANOVA comparing the two models. P-values and denominator degrees of freedom for these analyses were calculated using Satterthwaite’s approximations (Kuznetsova et al., 2013).

For the time-domain data, we identified time points where the experimental conditions were reliably different using a permutation based approach devised by Blair and Karniski (1993) and described in detail elsewhere (Appelbaum et al., 2006; Kohler et al., 2018). Briefly, this approach tests the null hypothesis that no differences were present between two conditions tested by making synthetic data sets in which the condition labels were randomly permuted across participants. 5000 permutations were randomly selected among the 32,768 that were possible for n = 15. For every permutation, we computed significances for a paired *t*-test comparing the waveforms and then calculated cluster-level statistics as the sum of t-values in the consecutive time points with *p*-values less than 0.05 (Maris and Oostenveld, 2007). We then took the maximum cluster-level statistic for each permutation, which constituted a non-parametric reference distribution of cluster-level statistics. We then rejected the null hypothesis if the cluster-level statistics for any consecutive sequence of significant *t*-scores in the original, non-permuted data was larger than 95% of the values in the null distribution. Because each permutation sample contributes only its maximum cluster-level statistic to the reference distribution, this procedure implicitly compensates for the problem of multiple comparisons, and is a valid test for the omnibus hypothesis of no difference between the waveforms at any time point. Furthermore, this test not only detects significant departures from the null hypothesis, but also localizes the time periods when such departures occur. However, since the correction procedure is tied to the length of the data and the somewhat arbitrary choice of keeping family-wise error at 5%, we therefore also present the uncorrected significance values (see red/yellow color maps in Figures 3, 5 and 7). By applying both statistical approaches, we are better able to identify time periods when the responses depart from the null hypothesis.

### Stimulus assessment

We directly assessed the images used in our experiments by generating images simulating ten trials worth of stimuli from each condition, and then independently estimating the number of dots, as well as four non-number dimensions. For each image, a mask was generated in which every non-background pixel was given the same value. Number of dots was then estimated as the number of contiguous regions in each mask, after first eroding the regions slightly to avoid counting abutting dots as a single region. The other parameters were similarly straightforward: The minimum number of pixels in the non-eroded contiguous region (“dot size”), the total number of non-background pixels (“total area”), convex hull of non-background pixels (“convex hull”), while “mean occupancy” was defined as convex hull divided by the number of dots (thus being an inverse measure of density). Together, these five factors provide a complete description of the variability in the images presented along the five dimensions.

### Early visual cortex simulation

Next, we wanted to estimate the degree to which early visual cortex might respond to the images we showed in the experiments by using a previously developed modeling approach (Van Rinsveld et al., 2020). The approach uses a cascaded, feedforward model of BOLD responses to visual stimuli, the second-order contrast (SOC) model, that first passes the image through a bank of contrast-normalized, localized, V1-like filters, and then reprocesses the output in a second stage that, among other things, measures the contrast variability in the output of the first stage (Kay et al., 2013). The SOC model has been fit to data for V1, V2, V3, and V4, and is effective at predicting responses to simple visual stimuli, including gratings and textures, while also capturing increased sensitivity to the structure of natural scenes in extra-striate visual areas (Kay et al., 2013). These features make it a reasonable model for estimating the activity that our dot images would elicit in early visual cortex. We applied the SOC model separately to each image in the ten-trial set described above, using model parameters for V1, V2, V3, and V4 provided in the original publication (Kay et al., 2013). We then summed the model output across each image to get the predicted average response of each brain area to each image update.

To summarize, the steps above produce a description of the stimulus along five different dimensions, as well as predicted brain responses across four early visual areas. To tie each of these variables to the brain responses measured in each of our experiments, and highlight the extent to which each changed systematically with reference and oddball updates, we computed the difference between each update and the preceding update, excluding the first update, separately for each variable. By plotting each difference estimates separately for oddball and non-oddball image updates, we can estimate the extent to which each variable would be expected to drive brain responses, which we can then compare to the predicted brain responses from the SOC model, and the measured brain responses in our three experiments.

## Results

### Experiment 1

In Experiment 1, the reference numerosity was 8 across all conditions, while the oddball could be either 5 or 8, with the latter serving as a control condition. To assess the temporal tuning of the approximate number sense, we recorded those two basic conditions with three distinct carrier and oddball frequency pairings: 6 Hz carrier with 1 Hz oddball, 3.75 Hz carrier with 0.75 Hz oddball and 3 Hz carrier with 0.5 Hz oddball.

The behavioral data (shown in Figure 2) indicated that performance was comparable for the two slower frequency pairs, with high *d*′ and very little bias. Performance was worse for 6 & 1 Hz, where participants exhibited a bias towards reporting that a number change had occurred.

**Figure 2:**
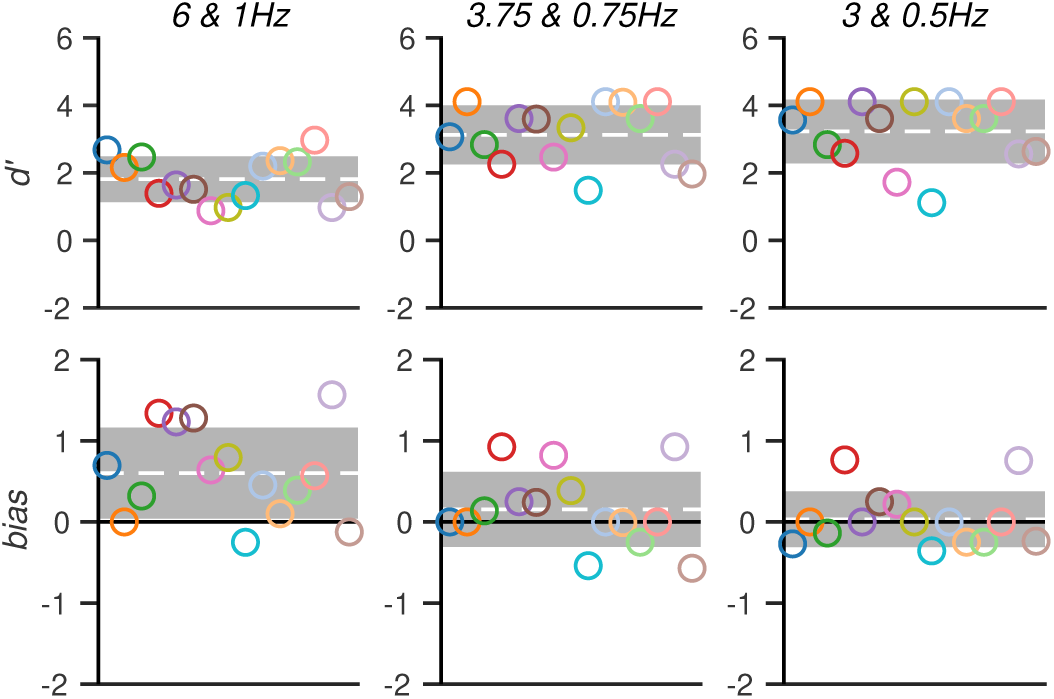
Behavioral Results from Experiment 1: Top half of the figure indicates d′ across the three different frequency pairs, while bias estimated as criterion (*c*) is shown on the bottom half. Each dot represents a participant, the gray region indicates the standard deviation, and the white dotted line indicates the averages.

The same SSVEP analysis was applied separately for each frequency pairing, which included deriving reliable components separately for each of the three pairs of stimulation frequencies and performing LMEAs for each of the first two components, separately for harmonics of the oddball and carrier frequency. The topography of the first component was centered over early occipital cortex across stimulation frequencies. By contrast, the second component was distributed more anteriorly and was centered on electrodes over parietal cortex (see Figure 2, middle column). The existence and magnitude of numerosity-related responses depended on both component and stimulation frequency. The first component exhibited no effects of numerosity for any of the frequency pairs. The oddball harmonics showed no main effects or interactions (6 & 1 Hz: smallest *p* = 0.369; 3.75 & 0.75 Hz: smallest *p* = 0.623; 3 & 0.5 Hz: smallest *p* = 0.390), while the carrier harmonics showed a main effect of harmonic (6 & 1 Hz: *F*(3, 98) = 5.647, *p* = 0.001; 3.75 & 0.75 Hz: *F*(3, 98) = 54.105, *p* < 0.0001; 3 & 0.5 Hz: *F*(3, 98) = 50.064, *p* < 0.0001), but no other main effects or interactions.

The second component had a different pattern of effects, and importantly showed numerosity-related responses for all three stimulation frequencies. The oddball harmonics showed main effects of numerosity (6 & 1: *F*(1, 98) = 5.091, *p* = 0.026; 3.75 & 0.75 Hz: *F*(1, 98) = 36.669, *p* < 0.0001; 3 & 0.5 Hz: *F*(1, 98) = 61.789, *p* < 0.0001). The oddball harmonics also showed main effects of harmonic for two of the three stimulation frequencies (3.75 & 0.75 Hz: *F*(3, 98) = 5.876, *p* < 0.001; 3 & 0.5 Hz: *F*(3, 98) = 4.259, *p* = 0.007). For 3 & 0.5 Hz, the interaction also reached significance (*F*(3, 98) = 4.149, *p* = 0.008) and interaction terms improved the model fit (*X*^*2*^(2) = 12.554, *p* = 0.006). The carrier harmonics showed main effect of harmonic (6 & 1 Hz: *F*(3, 98) = 11.063, *p* < 0.0001; 3.75 & 0.75 Hz: *F*(3, 98) = 2.982, *p* = 0.035; 3 & 0.5 Hz: *F*(3, 98) = 4.649, *p* = 0.004), but no other main effects or interactions.

We also projected the cycle-average waveforms through the component weights to visualize the responses in the time domain. Here all response frequencies contribute to the observed waveform. As expected from the frequency domain analysis, the waveform of the first component shows strong periodicity at the carrier frequency (Figure 3A, B, C, rightmost panels). Moreover, the control and odd-ball conditions (purple and green curves, respectively) have very similar waveforms, with only sporadic differences. This pattern of results is consistent with activity measured by this component being insensitive to numerosity.

**Figure 3:**
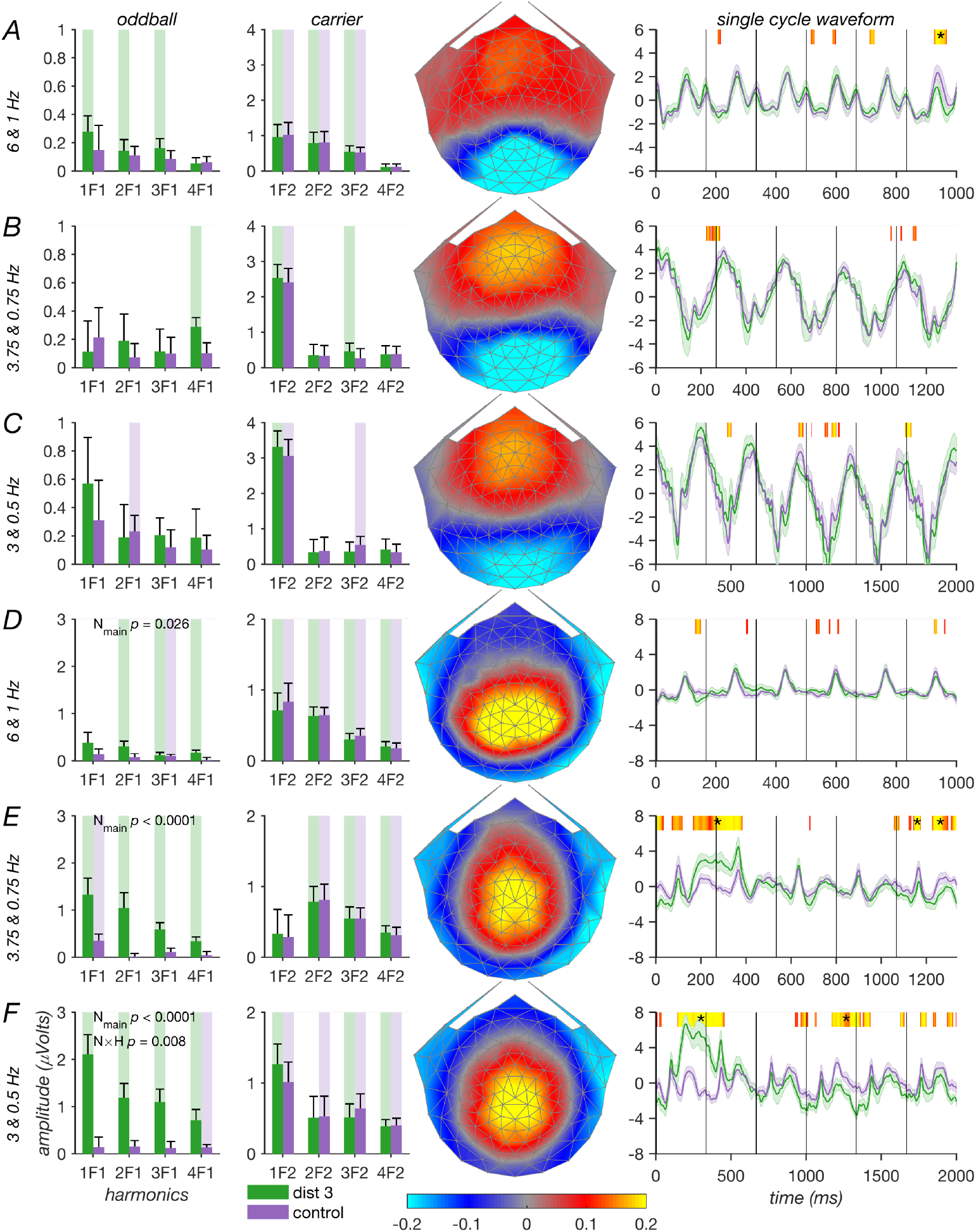
SSVEP results from Experiment 1: Shown for the first (A-C) and second (D-F) reliable components for each of the three frequency sets. Each row shows a separate component, with number oddball conditions plotted in green, while control conditions with no number oddballs are plotted in purple. The two leftmost columns indicate the first four harmonics of the oddball and carrier frequencies, respectively. Highlighted bars are significantly different from zero (*p* < 0.05). The topographies in the middle column indicate the topographies of the weights associated with each component. The rightmost column indicates the average response over a single cycle of the oddball frequency in the time domain. Orange-yellow patches indicate time points for which a paired t-test indicated that the number oddball and control conditions were significantly different (uncorrected *p* < 0.05), with more yellow indicating smaller p-values. Asterisks indicate segments of continuously significant time-points that permutation testing indicated were larger than what would be expected from chance (see text).

By contrast the waveform of the second component differentiates control and oddball conditions for lower-frequency pairs but not the higher frequency pair. Note the lack of significant runs for the 6 & 1 Hz data shown in Figure 3D, compared to the data for lower frequency pairs in Figure 3E and F where significant run-corrected differences between control and odd-ball conditions can be observed.

In summary, this experiment indicates that for all three stimulation frequencies, numerosity effects are seen for the second component that is distributed over parietal cortex, but not for the first component that is distributed over early occipital cortex. As we would expect from our experimental design, the numerosity effects are seen for the oddball harmonics, but not for the carrier harmonics that index responses to every image update. The numerosity effect is generally seen across multiple oddball harmonics contributing to differential activity between ~130 and 500 msec after the oddball appearance, but may be reduced for higher harmonics, as indicated by the significant interaction seen for 3 & 0.5 Hz. Finally, the numerosity effect is larger for the two slower stimulation frequencies, which implicates limitations in the temporal tuning of number processing mechanisms that are being measured. This is consistent with the diminished behavioral performance we observed for 6 & 1 Hz compared to the two slower frequency pairs.

### Experiment 2

In Experiment 2, we wanted to explore the effect of numerical distance between the reference and oddball numerosities. Because slower frequencies had produced the largest numerosity related signals in Experiment 1, we used 3 & 0.5 Hz stimulation frequencies for all conditions. The standard numerosity could either be 6 or 8, and each standard was paired with oddballs that were a distance of either 0, 1 or 3 away from the standard (6, 5 and 9 for reference 6; 8, 9 and 5 for reference 8). The behavioral data (shown in Figure 4) indicated that participants performed well on the task when the oddball was 3 away from the reference, but much worse when the distance was 1. Participants had a bias towards indicating that a number change had occurred when distance was 1 (most pronounced when a reference of 8 was paired with an oddball of 9), and a bias in the opposite direction when distance was 3.

**Figure 4:**
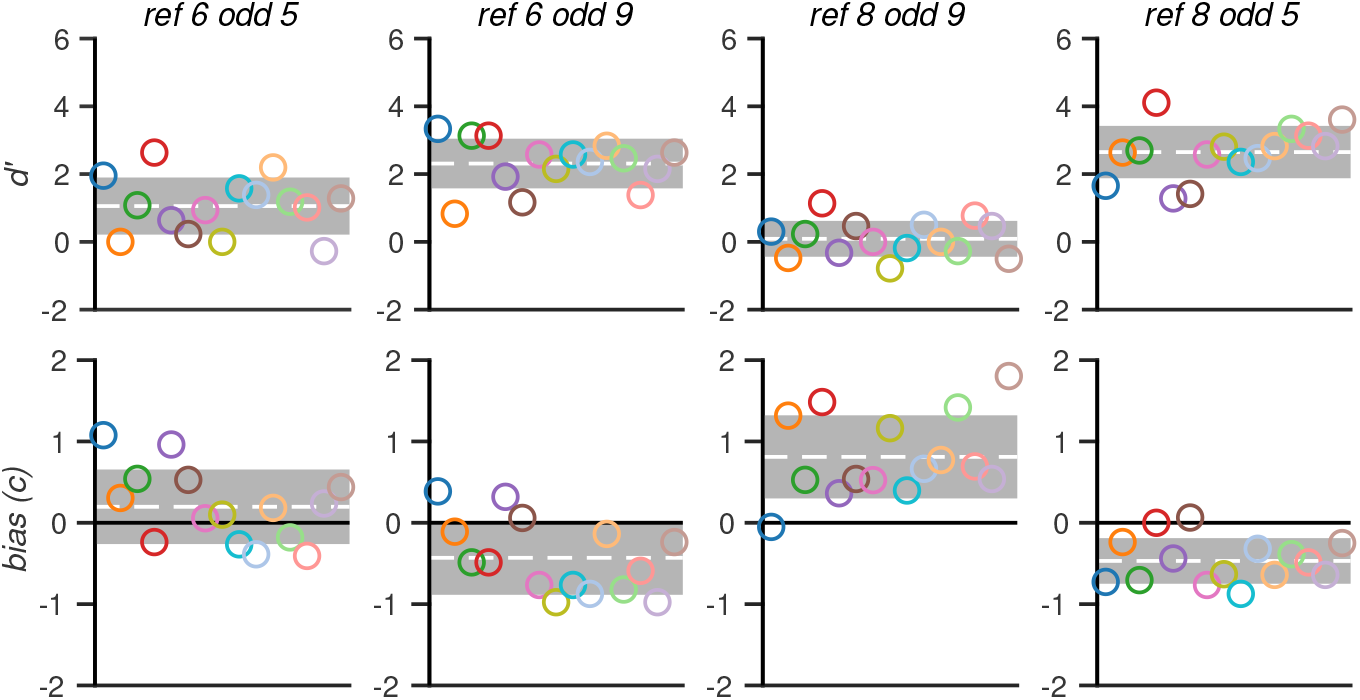
Behavioral Results from Experiment 2. d′ and bias estimated as criterion (*c*) across the four different reference and oddball pairs. Plotting conventions follow Figure 2, with each dot again represents a participant.

We derived reliable components based on all the SSVEP data, and then performed LMEAs for each of the first two components, separately for harmonics of the oddball and carrier frequency. The component topographies were similar to those observed in Experiment 1, with the first component centered over occipital cortex (see Figure 5E), and the second component distributed at electrodes over parietal cortex (see Figure 5F). As in the first experiment, the first component showed no evidence of numerosity responses, as evidenced by the absence of a main effect of numerosity in the oddball harmonics, which also failed to show any other main effects or interactions (smallest *p* = 0.278). The carrier harmonics showed a robust effect of harmonic regardless of reference numerosity (ref^6^: *F*(3, 154) = 59.003, *p* < 0.0001; ref^8^: *F*(3, 154) = 76.435, *p* < 0.0001), but no other main effects or interactions.

**Figure 5:**
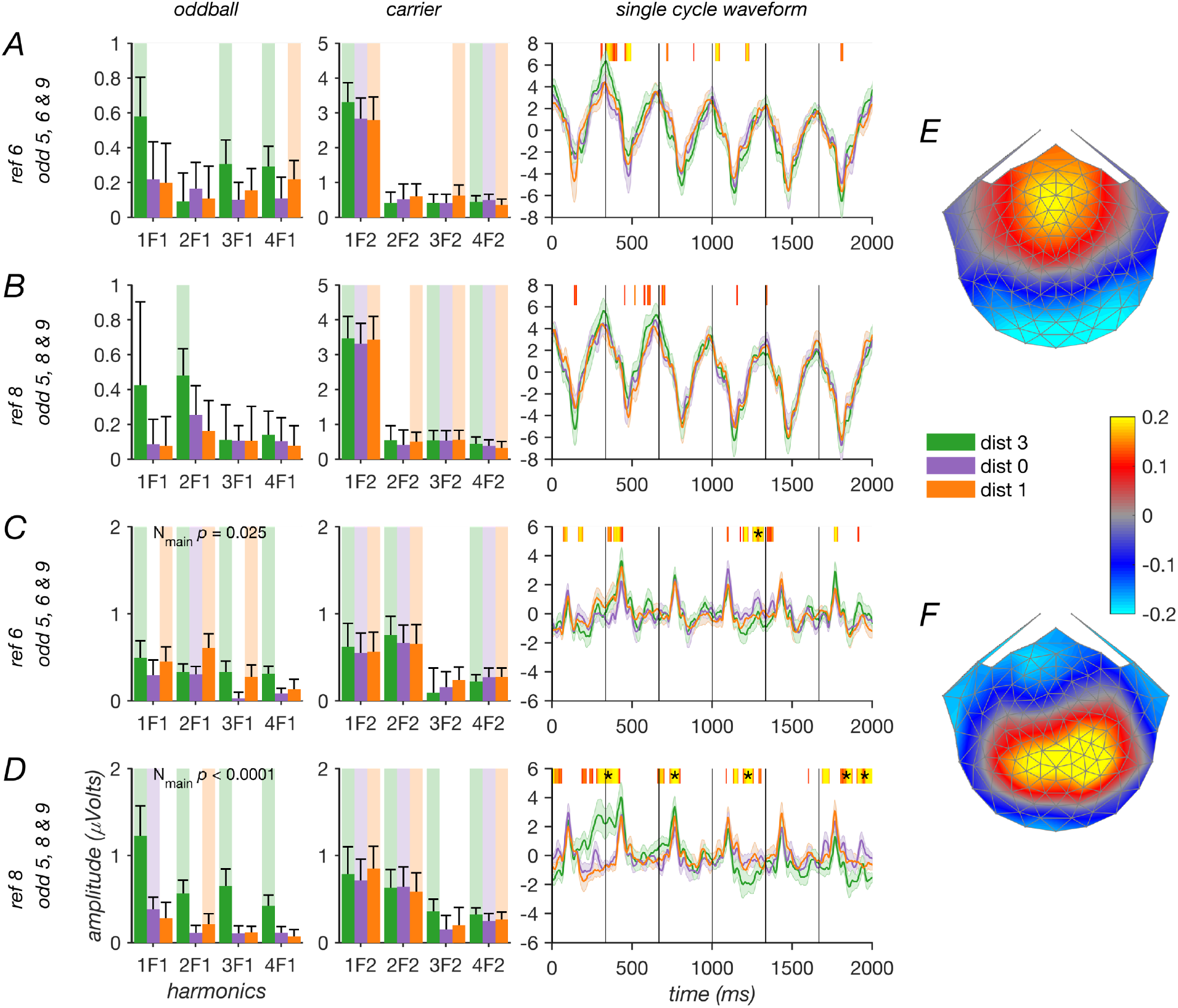
SSVEP Results from Experiment 2. Shown for the first (A-B) and second (C-D) reliable components. Each row shows a separate reference numerosity, with number oddball conditions with distance 3 plotted in green, distance 1 plotted in orange, and control conditions with no number oddballs are plotted in purple. The two leftmost columns indicate the first four harmonics of the oddball and carrier frequencies, respectively. Highlighted bars are significantly different from zero (*p* < 0.05). The topographies of the weights associated with the components are plotted in (E) for the first component and in (F) for the second. The rightmost column indicates the average response over a single cycle of the oddball frequency in the time domain. Orange-yellow patches indicate time points for which a paired t-test indicated that the oddball (distance = 3) and control (distance = 0) conditions were significantly different (uncorrected *p* < 0.05), with more yellow indicating smaller p-values. Note that distance = 1 was not included in this test. Asterisks indicate segments of continuously significant time-points that permutation testing indicated were larger than what would be expected from chance (see text).

The second component again showed evidence of a numerosity-related response, especially for the largest numerical distance. The oddball harmonics showed significant main effects of numerosity (ref^6^: *F*(2, 154) = 3.770, *p* = 0.025; ref^8^: *F*(2, 154) = 18.858, *p* < 0.0001) as well as main effects of harmonic (ref^6^: *F*(3, 154) = 3.798, *p* = 0.012; ref^8^: *F*(3, 154) = 5.120, *p* = 0.002), but no interactions. While main effects of numerosity were found for both conditions, the effects were most obvious for ref^8^, where the oddball with distance of 3 produced stronger responses than both the control condition and distance of 1 across the harmonics, consistent with the behavioral results. For ref^6^, oddball distances of 3 and 1 produced comparable responses, with both distances producing stronger responses than the control condition. This is somewhat consistent with the behavioral results, where distances of 3 and 1 produced more similar performances when the reference was six, than when the reference was 8. The carrier harmonics exhibited main effects of harmonic (ref^6^: *F*(3, 154) = 6.425, *p* < 0.001; ref^8^: *F*(3, 154) = 7.662, *p* < 0.0001), but no other main effects or interactions.

We again projected the bin-average time domain data through the component weights, and found that a numerical distance of 3 produced a robust differential response after run correction, replicating the numerosity-related response observed in the time-domain in Experiment 1.

### Experiment 3

In Experiment 3 we tested whether the somewhat weaker numerosity effect we observed for ref^6^ in Experiment 2 could be linked to the direction of the numerosity difference. In both Experiment 1 and 2, conditions with a reference of 8 and an oddball of 5 produced strong oddball responses, while the conditions in Experiment 2 with a reference of 6 and an oddball of 9 produced weaker responses. This led us to hypothesize that our method might be generally less sensitive to increases than to decreases in numerosity. To test this hypothesis, we again used the same stimulation frequencies for all conditions, 3 & 0.5 Hz. The reference numerosity could either be 5, 6, 8 or 9, and each reference was paired with oddballs that were either 0 or 3 away from the reference (ref^5^: 5 or 8; ref^6^: 6 or 9; ref^8^: 5 or 8; ref^9^: 6 or 9). The behavioral data (shown in Figure 6) again showed that participants performed well on the task, with perhaps a slight decrease in performance for higher numerosities. Participants had very little bias regardless of reference numerosity.

**Figure 6:**
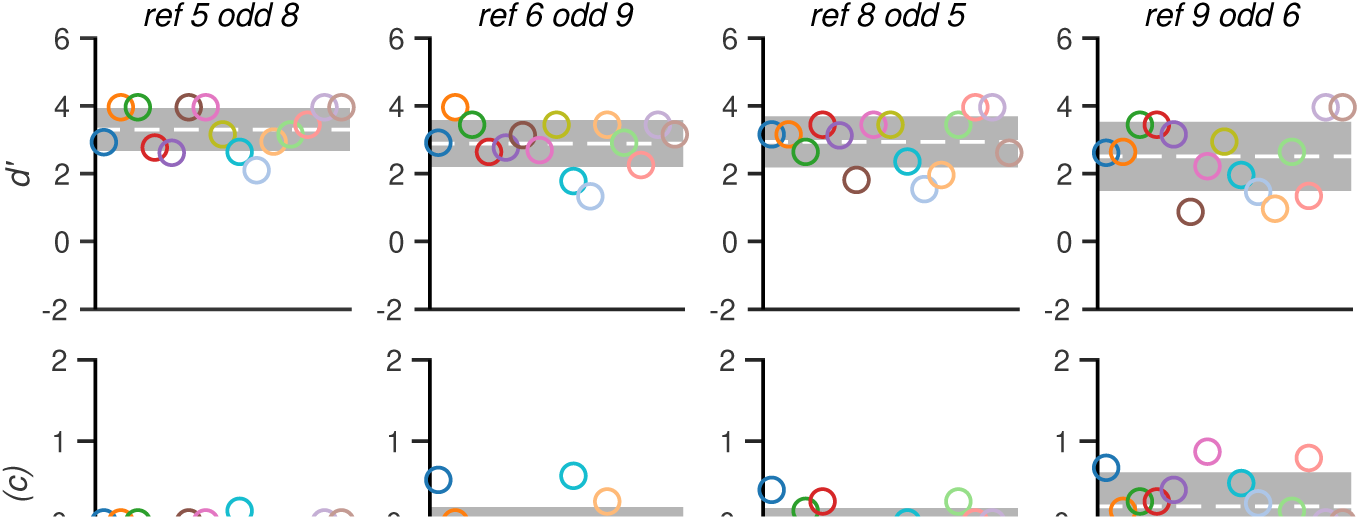
Behavioral Results from Experiment 3: d′ and bias estimated as criterion (*c*) across the four different reference and oddball pairs. Plotting conventions follow Figure 2, with each dot again represents a participant.

We again derived reliable components based on all the SSVEP data, and then performed LMEAs for each of the first two components, separately for harmonics of the oddball and carrier frequency. As in the first two experiments, the first component was centered over occipital cortex (see Figure 7I), while the second component was centered over parietal cortex (see Figure 7J).

**Figure 7:**
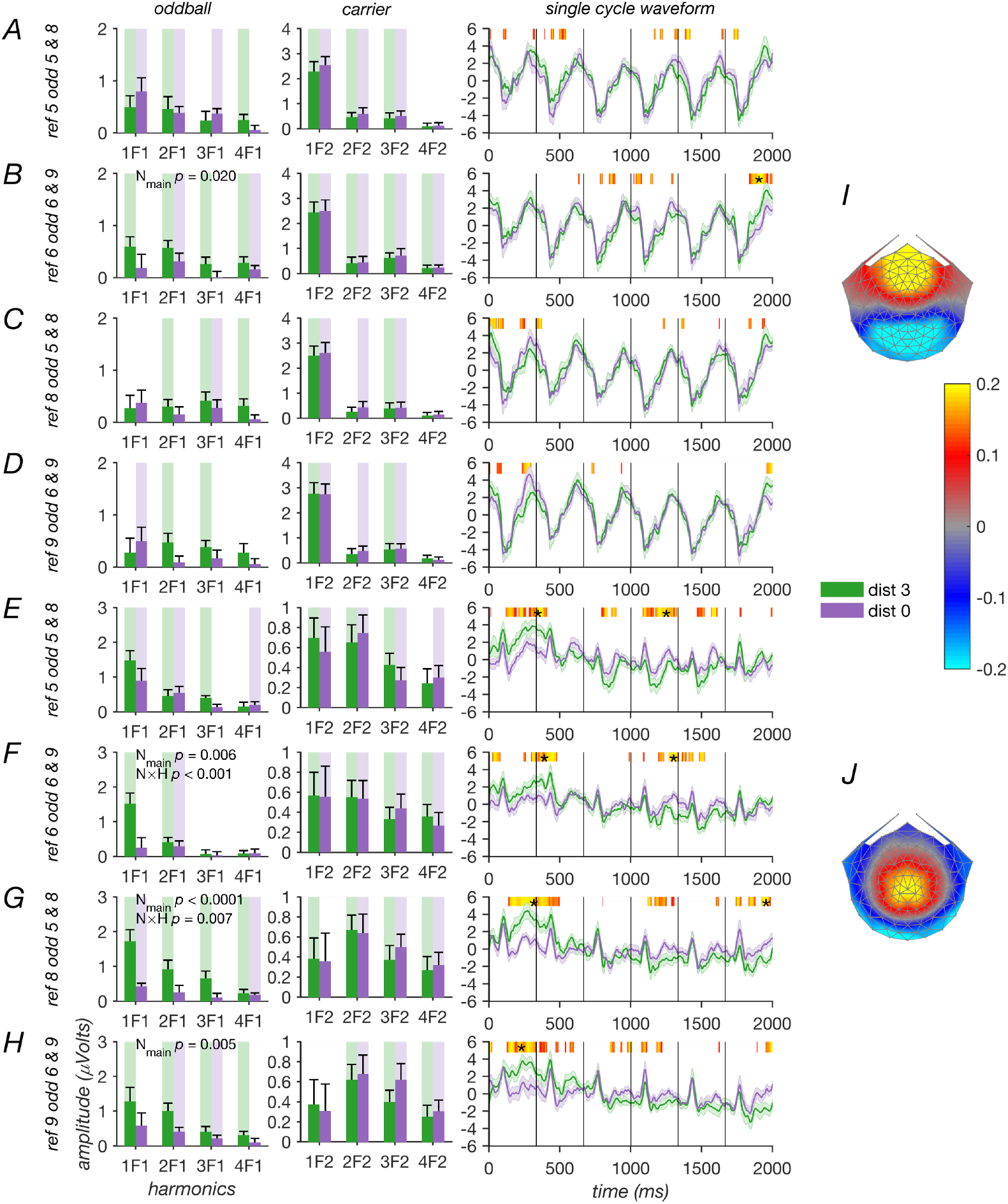
SSVEP results from Experiment 3. Shown for the first (A-D) and second (E-H) reliable components. Each row shows a separate reference numerosity, with number oddball conditions with distance 3 plotted in green and control conditions with no number oddballs plotted in purple. The two leftmost columns indicate the first four harmonics of the oddball and carrier frequencies, respectively. Highlighted bars are significantly different from zero (*p* < 0.05). The topographies of the weights associated with the components are plotted in (I) for the first component and in (J) for the second. The rightmost column indicates the average response over a single cycle of the oddball frequency in the time domain. Orange-yellow patches indicate time points for which a paired t-test indicated that the number oddball and control conditions were significantly different (uncorrected *p* < 0.05), with more yellow indicating smaller p-values. Asterisks indicate segments of continuously significant time-points that permutation testing indicated were larger than what would be expected from chance (see text).

Unlike the previous experiments, we saw weak evidence of numerosity responses in the first component. for 1 of our 4 reference numerosities. The oddball harmonics for ref^6^, exhibited a main effect of numerosity (*F*(1, 112) = 5.592, *p* = 0.020), with no other main effects of interactions. Ref^5^ showed a main effect of harmonic (*F*(3, 98) = 3.233, *p* = 0.026), but the three other references produced no other main effects or interactions (smallest *p* = 0.262). The carrier harmonics showed strong main effects of harmonic for all 4 references (ref^5^: *F*(3,98) = 43.972, *p* < 0.0001; ref^6^: *F*(3,98) = 35.427, *p* < 0.0001; ref^8^: *F*(3,98) = 49.733, *p* < 0.0001; ref^9^: *F*(3,98) = 52.615, *p* < 0.0001), but no other main effects or interactions.

For the second component, the oddball harmonic responses showed significant main effects of numerosity for 3 of 4 references (ref^5^: *F*(1, 98) = 2.160, *p* = 0.145; ref^6^: *F*(1, 98) = 7.882, *p* = 0.006; ref^8^: *F*(1, 98) = 26.543, *p* < 0.00001; ref^9^: *F*(1, 98) = 8.371, *p* = 0.005). All four reference numerosities also produced significant main effects of harmonic (smallest *p* = 0.002), and two produced significant interactions (ref^5^: *F*(3, 98) = 1.684, *p* = 0.176; ref^6^: *F*(3, 98) = 5.946, *p* < 0.001; ref^8^: *F*(3, 98) = 4.310, *p* = 0.007; ref^9^: *F*(3, 98) = 0.823, *p* = 0.484). The carrier harmonics showed significant main effects of harmonic for 2 out of 4 reference numerosities, with ref^8^ also approaching significance (ref^5^: *F*(3, 98) = 4.809, *p* = 0.004; ref^6^: *F*(3, 98) = 1.115, *p* = 0.347; ref^8^: *F*(3, 98) = 2.502, *p* = 0.064; ref^9^: *F*(3, 98) = 3.240, *p* = 0.025), but no other main effects of interactions. It is perhaps surprising that ref^5^ did not produce a significant main effect of numerosity or an interaction, but we note that for at least two of the four harmonics tested, the effect was in the right direction. Furthermore, when we projected the cycle-average time domain through the component weights, the difference in response waveforms between control and oddball conditions was of similar magnitude across all four reference numerosities. Based on this evidence, we conclude that the direction of the numerosity change is not an important determinant of the oddball response magnitude.

In summary, the results from all three experiments showed a clear numerosity-oddball response that was component and stimulation frequency specific. Numerosity responses were almost exclusively seen for components centered over parietal cortex, with ref^6^ in Experiment 3 (1 out of 9 comparisons) being the only exception where a numerosity effect is observed over occipital cortex. Numerosity responses were strongest for slower stimulation frequencies (0.75 & 3.75 Hz and 0.5 & 3 Hz) and could be produced for both decreases and increases in numerosity, although the latter effect may be less consistent.

### Stimulus Assessment

We ran two distinct analyses on sample images from the conditions used in the three experiments. The goal of the first analysis was to provide a complete and independent description of the variability in the images presented along the five dimensions of variation used to create the dot clouds.

We estimated sample statistics for dot size, total area, convex hull, and density, as well as numerosity based on 10 trials worth of test images, as described in the Methods. The distribution of changes along a dimension on each image was overall quite similar between carrier and oddball updates for each of the dimensions of variation, and any differences we did observe between oddball and carrier updates were well-matched by differences in the associated control condition (see Figure 8). The results thus confirm that we were successful in generating control conditions that elicited no change in numerosity yet were well-matched on other dimensions to the conditions for which the oddball was associated with a change in numerosity.

**Figure 8:**
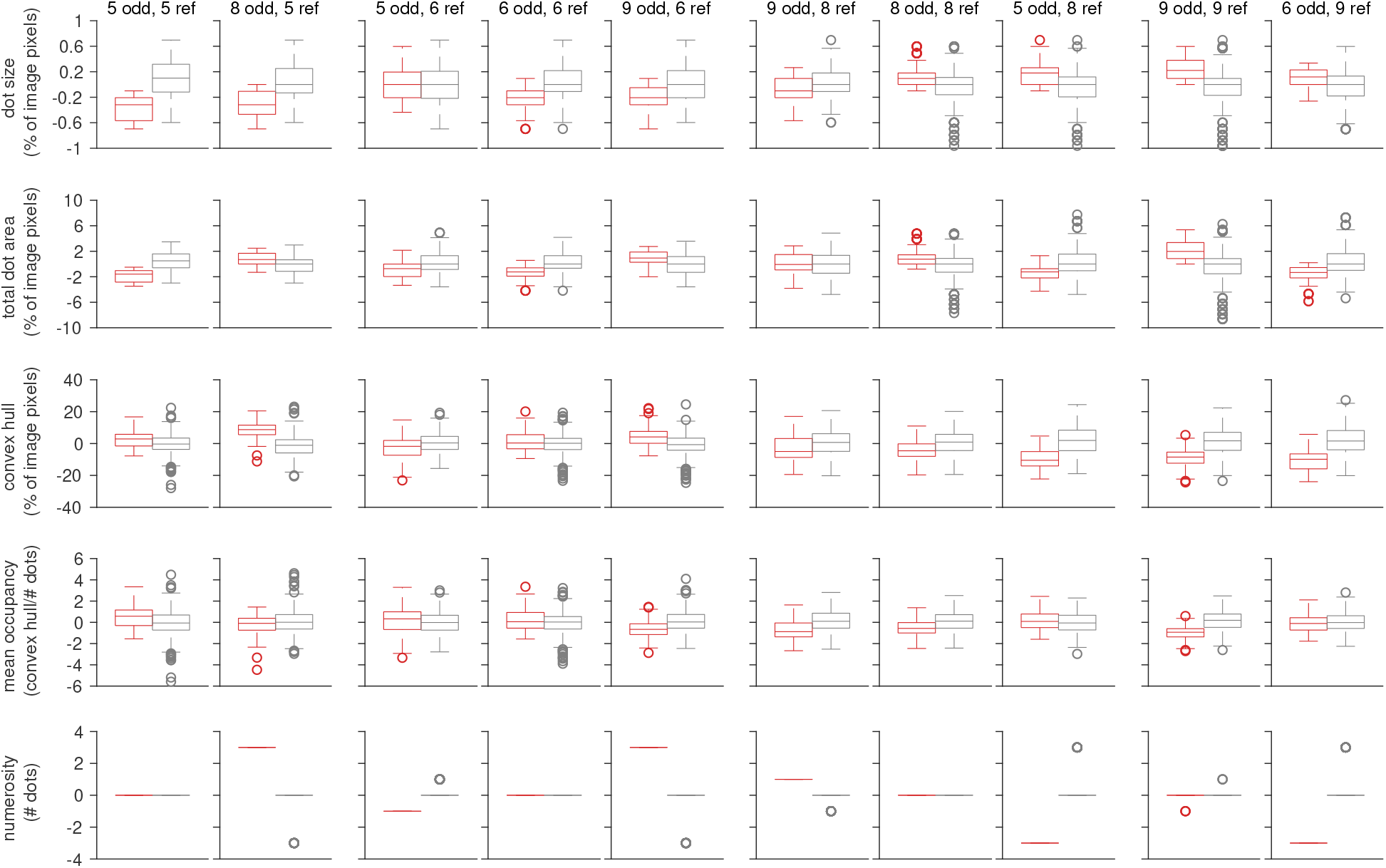
Change on oddball and reference image updates for the stimuli used in all three experiments. Each column represents a condition, as indicated at the top of the plot. Note that columns are grouped together by reference numerosity. Each row indicates the difference from the previous image update on five distinct stimulus dimensions. Reference and oddball updates are plotted in gray and red, respectively. The dimensions were computed as described in the text. Note that different plots use different units, as indicated on the y-axis. For each boxplot, the central mark is the median, the edges of the box are the 25th and 75th percentiles, the whiskers extend to the most extreme data points not considered outliers, and the outliers indicated with dots.

The goal of the second analysis was to test whether a simple model of early visual cortex, designed to capture activity in visual cortex elicited by simple gratings, textures and natural scenes, and fit to functional MRI data could explain the results we observed. To that end, we applied to the images used in the experiments to the Second-Order Contrast (SOC) model (Kay et al., 2013; Van Rinsveld et al., 2020). The model output was well-aligned with our SSVEP data, in that oddball stimuli with a numerical distance of 3 produced model outputs that were larger than those produced by both oddball stimuli with a numerical distance of 1 and carrier updates (Figure 9).

**Figure 9:**
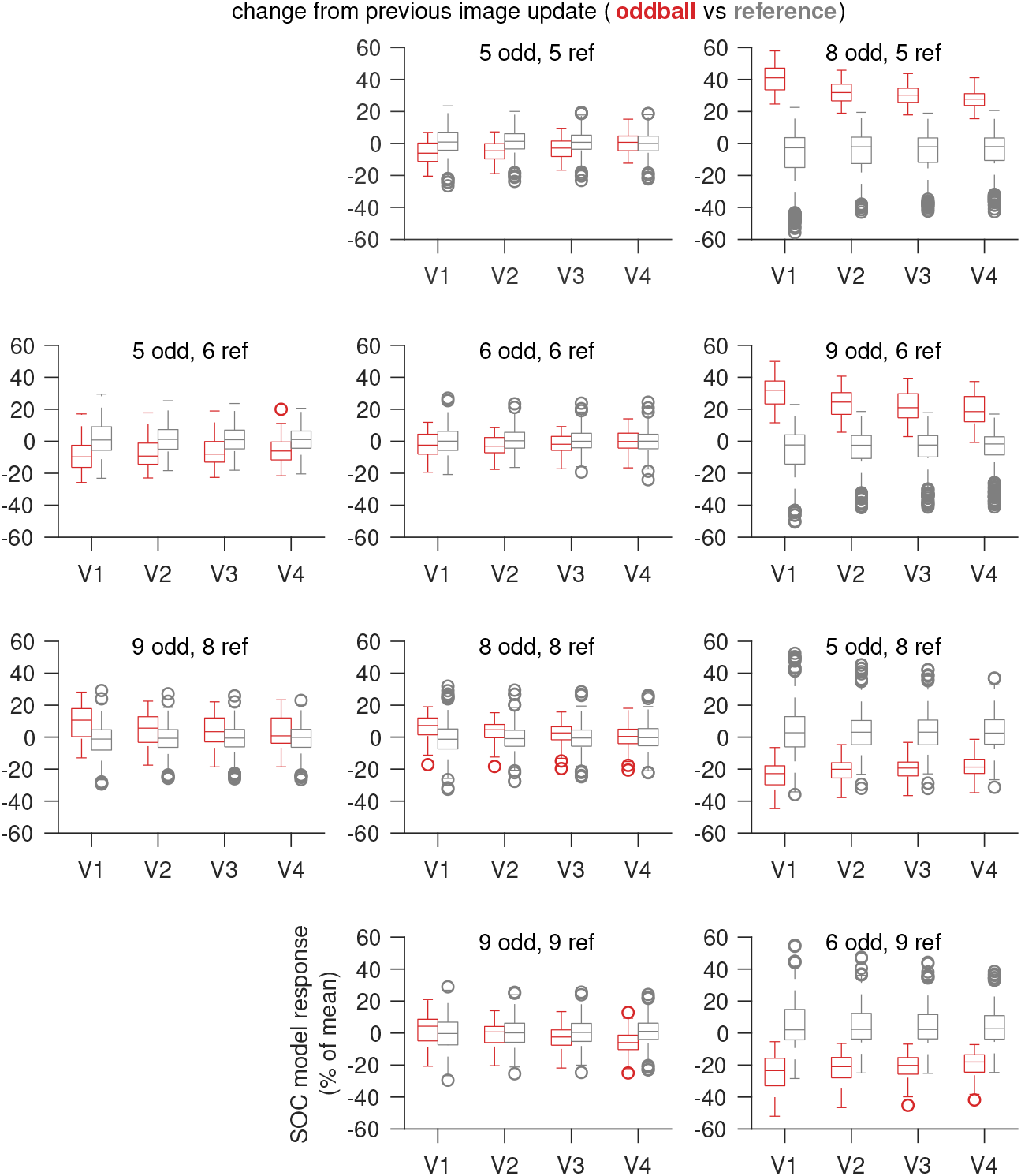
Simulated responses for stimuli used in the 3 experiments. Change in SOC model responses from previous image update, plotted separately for oddball (in red) and reference (in gray) updates, as boxplots with the bottom and top edges of the box indicating the 25th and 75th percentiles, with whiskers extending out to 1.5 × the interquartile range, and anything beyond that considered outliers. The outliers are plotted with circular markers. Experiment 1 used an 8-dot reference stimulus and a 5-dot oddball, with an 8-dot reference and 8-dot oddball as control (top row). Experiment 2 used the 6 different stimuli shown in the two middle rows. Experiment 3 used the 8 different stimuli in the two rightmost columns.

Overall, these results suggest that computations that have been associated with feature processing in visual cortex are able to signal numerosity in a fashion that is semi-quantitatively similar to our measured responses.

## Discussion

By manipulating the temporal presentation rate of dot clouds containing different numbers of dots, we have shown that both behavioral discrimination and evoked activity over parietal electrodes is degraded at higher presentation rates. Previous high presentation rate SSVEP studies have found number-related responses on electrodes over early visual cortex in experiments where numerosity was task-irrelevant (Guillaume et al., 2018; Park, 2018; Van Rinsveld et al., 2020). Our results suggest that the faster stimulation rates used in these studies may have exceeded the temporal limit of parietal number mechanisms responsible for conscious report of numerosity.

A current debate in the numerical cognition literature centers around whether number is a primary stimulus attribute like orientation or color, encoded in early cortical areas or rather is derived from more abstract processing in higher level cortex. These views are not mutually inconsistent, and our results provide evidence for both. Our parietally focused second component is consistent with the many reports of number representations in higher-level, notably parietal cortex (Lasne et al., 2019). Importantly the magnitude of our second component scales with numerical distance as has been reported in fMRI studies of parietal areas (Lasne et al., 2019).

Our analysis of five stimulus dimensions (dot size, total dot area, convex hull, mean occupancy and numerosity) in our stimulus set, shows that oddball updates are not associated with systematic change in the non-number dimensions. In line with previous work using similar stimuli (Van Rinsveld et al., 2020), this suggests that the numerosity oddball responses we measure are not trivial consequences of stimulus confound.

Our simulation results and related results from previous work (Van Rinsveld et al., 2020) suggest that numerosity-related signals could first arise through the relatively simple computations available in early visual cortex. The SOC model used here and in the previous studies instantiates canonical computations such as spatial filtering, exponentiation and normalization that are common throughout cortex (Carandini and Heeger, 2011). However, unlike several previous studies (Guillaume et al., 2018; Park, 2018; Lucero et al., 2020; Van Rinsveld et al., 2020), we find only weak evidence of a number response in our occipitally focused RC1. For 8 of our 9 comparisons across our three experiments, the occipital component failed to produce a significant effect of numerosity. Our stimulation conditions differ from those of previous studies in the lower stimulation rates we used, as well as in the use of luminance-balanced dots. The lower stimulation rate may not produce sufficiently robust responses in early visual areas for us to measure them reliably, while these rates may be more appropriate for driving robust responses in higher level cortex. It is also possible that better control of luminance-related responses through the use of balanced dots could reduce occipital responses. While the modeling results suggest a basis for number effects attributable to early visual cortex (Park et al., 2016; Fornaciai et al., 2017; DeWind et al., 2019), they are not sufficient to explain our results from parietal electrodes.

### The role of higher-level cortical areas

Studies using transient ERPs have demonstrated number-related visual responses over parietal electrodes that were sensitive to numerical difference during the performance of non-symbolic comparison tasks (Libertus et al., 2007; Heine et al., 2011; Heine et al., 2013). The latency of this differential activity was around 180 msec in adults (Libertus et al., 2007) or 300-600 msec in children (Heine et al., 2011; Heine et al., 2013). The slower stimulation rates for the oddball in our experiments are in some sense analogous to the widely separated presentations typically used in ERP designs and this may be a factor in the similarity of the response topography.

Two studies have explicitly compared number related activations in early visual areas to activations found in parietal cortex. One study using well controlled stimuli has reported number-related activity in early visual cortex, but not in parietal cortex (DeWind et al., 2019). In that study, numerosity was not task relevant. Another study found that while numerosity could be decoded from voxel activity in early visual cortex, individual variation in activation in these areas was not predictive of an explicit numerical judgment task (Lasne et al., 2019). Rather, behavioral Weber fractions were predictable from parietal areas, especially in right lateral intraparietal cortex. The correlational analysis performed in that study had low statistical power (n=12), so the null result from early visual cortex may not be in direct conflict with the SSVEP results implicating early visual area sensitivity in determining sensitivity at higher levels of cortex (Guillaume et al., 2018).

With respect to the importance of higher-level cortex in conscious number processing, our second component which is centered on electrodes over parietal cortex and scales with numerical distance is suggestive. This component, like behavioral discrimination of numerosity, is adversely affected by high temporal rates of presentation. The temporal limitation we report here for numerosity is reminiscent of the temporal limitations previously reported for a number of higher-order visual processes (Holcombe, 2009). For example, the temporal resolution of luminance detection is greater than 50 Hz (de Lange, 1958; Kelly, 1959), while the temporal limit for visual binding of words is 5Hz or less (Holcombe and Judson, 2007), a limit similar to the limit on behavior we see here. Approximate number representations may take two forms in the brain, each with different temporal limits. The first form could arise from implicit, domain general visual processing mechanisms that have a relatively high temporal resolution (at least 10 Hz) and reside in early visual areas (Guillaume et al., 2018; Park, 2018; Van Rinsveld et al., 2020). The output of this level of processing may then be integrated in higher level cortical areas where it is made available for conscious access and use by other cognitive systems involving learning and reasoning.

## Acknowledgements

PJK acknowledges funding support from the Canada First Research Excellence Fund and the National Sciences and Engineering Research Council of Canada.

